# Non-classical estrogen signaling inhibits melanoma and improves response to PD-1 blockade

**DOI:** 10.1101/146498

**Authors:** Christopher A. Natale, Jinyang Li, Junqian Zhang, Ankit Dahal, Ben Z. Stanger, Todd W. Ridky

**Affiliations:** Perelman School of Medicine, Department of Dermatology, University of Pennsylvania, Philadelphia, PA 19104, USA; Abramson Family Cancer Research Institute, Perelman School of Medicine, University of Pennsylvania, Philadelphia, Pennsylvania 19104, USA.

## Abstract

Female sex and pregnancy are associated with reduced risk of melanoma and improved stage specific survival; however, the mechanism underlying this apparent clinical benefit is unknown. We previously discovered that pregnancy-associated 17β-estradiol drives melanocyte differentiation by activating the nonclassical G-protein coupled estrogen receptor (GPER). Here, we show that pregnancy inhibits melanoma, and that transient GPER activation induces long-term changes in melanocytes, which are associated with increased cellular differentiation and resistance to melanoma. A selective GPER agonist induced c-Myc protein degradation, slowed tumor growth, and inhibited expression of immune suppressive proteins including PD-L1, suggesting that GPER signaling may render melanoma cells more vulnerable to immunotherapy. Systemically delivered GPER agonist was well tolerated, and cooperated synergistically with PD-1 blockade in melanoma-bearing mice to dramatically extend survival. These results thus define GPER as a target for differentiation-based melanoma therapy.

**Significance:** Immune checkpoint inhibitors including αPD-1 induce durable remissions in only 30% of patients with advanced melanoma. This work demonstrates that αPD-1 efficacy is significantly improved by systemic delivery of a selective agonist of the nonclassical estrogen receptor, GPER, which drives melanoma differentiation and renders tumors more vulnerable to the immune system.

## Introduction

Melanoma is the most deadly form of skin cancer and incidence is rising worldwide. Despite recent advancements in immunotherapies, the majority of patients with metastatic melanoma still succumb to their disease and mean survival is 23 months (1, 2). There is an acute need for new therapeutic strategies that augment the efficacy of standard-of-care immune checkpoint inhibitors. Clues to potential new therapeutic targets for melanoma may be found in 50 year old observations (3), validated in recent studies, that female gender, history of multiple pregnancies, and decreased maternal age at first birth are each associated with decreased melanoma incidence and favorable prognosis (4–7). Although the mechanism of this protective effect is unknown, the clinical association suggests that sex hormone signaling is involved. We hypothesized that understanding the relevant hormones, receptors, and downstream signaling events activated in melanocytes by pregnancy-associated sex steroids would help define the mechanism of the female melanoma protective effects and suggest new therapeutic opportunities.

In previous studies we determined that estrogen, which is higher in females, especially during pregnancy, acts directly on skin melanocytes to drive melanocyte differentiation and pigment production (8). This was a somewhat unexpected result, as melanocyte differentiation was previously thought to regulated primarily by the activity of the melanocortin receptor 1 (MC1R), a stimulatory Gs-coupled G protein-coupled receptor that signals through adenylate cyclase, cAMP, and PKA to activate CREB. CREB then drives expression of microphthalmia transcription factor (MITF), the master regulator of melanocyte differentiation (9–11). Consistent with the premise that canonical GPCR signaling regulates melanocyte differentiation state, we found that the estrogen effects in melanocytes are mediated entirely through a G protein-coupled receptor, named G-protein coupled estrogen receptor (GPER), which was not previously known to have activity in melanocytes. GPER activates signaling pathways that are completely distinct from classical estrogen receptors (12), but that converge with MC1R signaling at the level of adenylate cyclase. Although there are no approved drugs that specifically target GPER, we determined that GPER is activated in both female and male melanocytes by estrogen, as well as by a selective agonist (G-1) that activates GPER signaling without affecting the activity of classical estrogen receptors (ERα/β). (13). Here we show that GPER activation in melanoma cells induces a constellation of durable phenotypic changes that inhibit tumor growth, and also render tumor cells more susceptible to clearance by native immune cells, which increases the clinical efficacy of anti-PD-1 immune checkpoint blockade. Selective GPER agonists, may represent an entirely new class of differentiation-promoting anti-cancer agents.

## Results

To test whether pregnancy affects melanoma development, we used genetically-defined human melanoma (heMel) xenografts (14, 15). In this tissue model, primary human melanocytes were engineered with lentiviruses to express mutant oncoproteins commonly associated with spontaneous human melanoma (14) including BRAF^V600E^ (doxycycline-inducible), dominant-negative p53^R248W^, active CDK4^R24C^ and hTERT (Supplementary Fig. S1A). The oncogene expressing melanocytes were combined with primary human keratinocytes and native human dermis to construct functional 3-dimensional human skin tissues that were grafted into the orthotopic location on the backs of female mice (Supplementary Fig. S1B). After grafts healed, mice were randomized and separated into nonbreeding or breeding groups (Fig. 1A). Doxycycline chow was then provided to induce the BRAF^V600E^ oncogene in all animals. After 15 weeks and 3 consecutive pregnancies in the breeding group (or no pregnancies in the nonbreeding group), human tissues were harvested and analyzed histologically. Grafts from the nonbreeding group developed into melanocytic neoplasms with hallmark features of human melanoma including large, mitotically active melanocytic nests with cellular atypia (Fig. 1B-D and Supplementary Fig. S1C). In contrast, tissues from the breeding group were relatively unremarkable, and contained primarily quiescent, single, non-proliferating melanocytes that were confined to the basal epidermal layer. These results show that repeated pregnancies inhibit the growth of BRaf-driven human melanocytic neoplasia.

**Figure 1.**
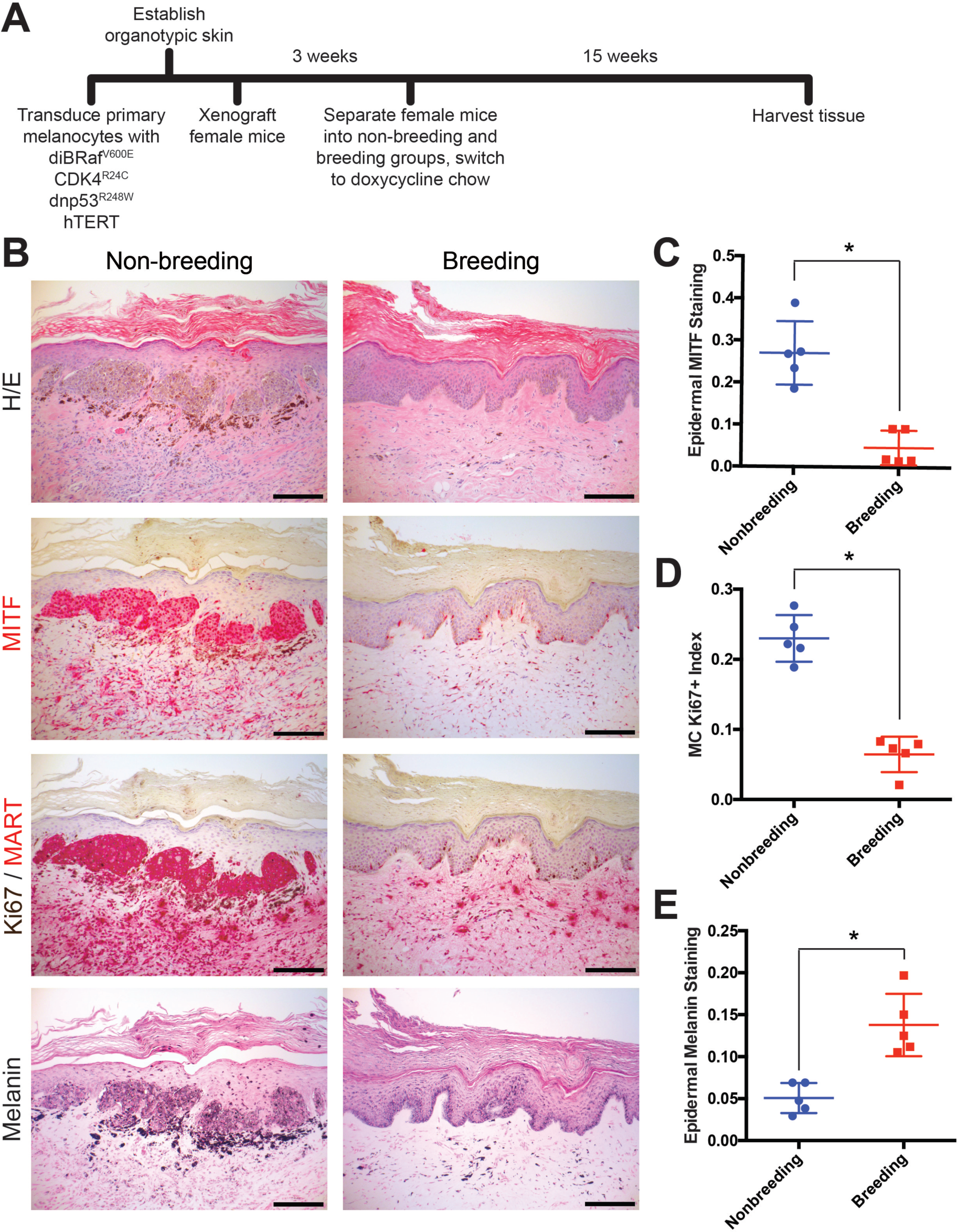
Multiple pregnancies inhibit melanomagenesis. **A**, Experimental timeline of genetically-defined human xenograft melanoma on SCID mice, n = 5 per group. **B**, Histologic characterization of representative orthotopic skin and resulting tumors, including hematoxylin and eosin (H/E), melanocyte and proliferation markers MITF, Ki67/MART, and Fontana Masson (Melanin). Scale bars = 100µM. **C-E**, Quantification of epidermal MITF staining (**C**), Ki67 proliferation index (**D**) and melanin staining in epidermal keratinocytes (**E**), * denotes significance by the Mann-Whitney test.

The primary role of a fully differentiated epidermal melanocyte is to produce melanin pigment that protects the skin from ultraviolent radiation (11, 16, 17). As with most cell types, melanocyte differentiation and proliferation are inversely correlated, and melanocytes in normal skin rarely proliferate outside of cycling hair follicles (18, 19). Melanoma tissue is generally less differentiated than normal melanocytes or benign nevi. In our xenograft studies, pregnancy was associated with an increase in melanocyte differentiation compared to the nonbreeding group, as evidenced by the relative lack of proliferating melanocytes and corresponding increase in epidermal melanin. Although the nonbreeding group, which developed melanomas, had significantly more melanocytes in the grafted skin than the breeding group, melanin abundance within the surrounding epidermal keratinocytes was dramatically reduced (Fig. 1E). These data suggest that pregnancy inhibits melanoma development by inducing melanocyte differentiation.

To test whether pregnancy-associated hormones induce long-lasting changes in melanocyte differentiation, we transiently exposed primary human melanocytes to estrogen or progesterone. Consistent with our previous studies where we discovered that continues exposure to estrogen and progesterone reciprocally regulate melanocyte pigmentation and differentiation state through the stimulatory GPCR, GPER, or inhibitory GPCR PAQR7, respectively (8), estrogen drove differentiation associated with increased melanin production, while progesterone had opposite effects (Fig. 2A). After hormone withdrawal, progesterone treated cells quickly returned to their baseline level of melanin production. In contrast, estrogen treated cells remained more differentiated after estrogen withdrawal, and stably produced more melanin through continual cell divisions over the subsequent 50 days. A subset of cells differentiated by transient exposure to estrogen was subsequently treated with progesterone. This reversed the estrogen effects, and melanin production decreased to the sub-baseline level seen upon initial progesterone treatment. Remarkably, after progesterone withdrawal, these cells fully returned back to the heightened differentiation state induced by the initial estrogen exposure (Fig. 2A). Consistent with increased cellular differentiation, estrogen exposure was associated with stable increases in classic melanocyte differentiation antigens including tyrosinase and MC1R (Fig. 2B). These results indicate that estrogen signaling, even transiently, induces a durable, long-lasting differentiation program in melanocytes.

**Figure 2.**
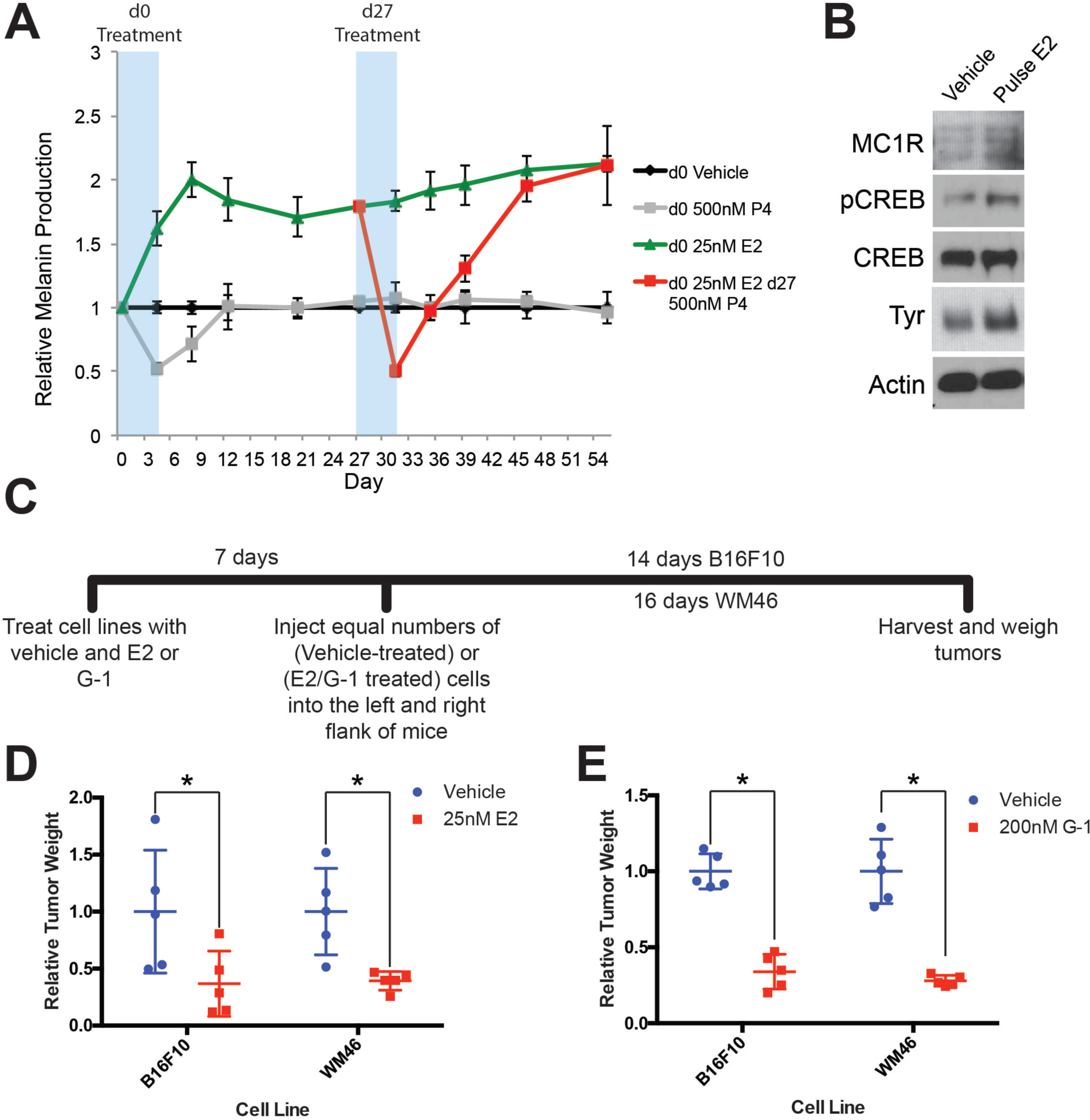
GPER signaling drives stable differentiation in normal human melanocytes and in melanoma. **A**, Long-term melanin assay in which normal human melanocytes were transiently treated with progesterone (P4), or estrogen (E2). Subsets of these groups (Red) were treated with an additional transient pulse of P4 at Day 27. Error bars equal the standard deviation of the samples. **B**, Western blot of melanocyte differentiation markers after a transient, 4-day treatment with either vehicle or estrogen, followed by an 8 day withdraw period. **C**, Experimental timeline of estrogen or GPER agonist (G-1) pre-treatment of mouse and human melanoma cells, n=5 per group. **D**, Relative tumor weights of mouse and human melanomas pre-treated with estrogen, * denotes significance by the Mann-Whitney test. **E**, Relative tumor weights of mouse and human melanomas pre-treated with G-1, * denotes significance by the Mann-Whitney test.

To determine whether estrogen drives differentiation in melanoma, we treated mouse (B16F10) or several human melanoma cells (WM46, WM51, WM3702) with either estrogen, or the specific GPER agonist G-1, which has no activity on the classic nuclear estrogen receptor (13). Consistent with changes observed in heMel cells, estrogen or G-1 decreased tumor cell proliferation and increased melanin production, independent of the specific oncodrivers (BRAF^V600E^ or NRas^Q61L^) in these melanoma cells (Supplementary Fig. S2A-D). To test whether transient GPER signaling induces a persistent differentiation state in melanoma cells that affects subsequent tumor growth in vivo, we treated melanoma cells with estrogen, G-1, or vehicle in vitro, and subsequently injected equal numbers of treated cells into host mice (Fig. 2C). Pretreatment with estrogen or G-1 markedly reduced subsequent tumor size (Fig. 2D-E), indicating that transient GPER activation has durable effects on melanoma cells that limit tumor growth in vivo.

Amplification of *c-Myc* – a transcription factor that antagonizes differentiation and promotes proliferation, survival, and escape from immune surveillance – is one of the most common genetic alterations in human cancers, including melanoma (20, 21). We found that GPER signaling in melanoma cells stably depleted c-Myc protein, and induced a relative growth arrest. This was associated with persistent hypophosphorylation of Rb, increased expression of HLA, and reduced expression of PD-L1 (Fig. 3A-D and Supplementary Fig. S2E-F). c-Myc loss is a major mediator of the anti-proliferative effects of GPER signaling, as melanoma cells engineered to maintain c-Myc protein in the face of GPER activation were resistant to G-1. (Fig. 3E). To verify that G-1 effects in melanoma were mediated entirely through GPER, we utilized a selective GPER antagonist, G-36 (22), that specifically inhibits GPER, but that does not bind ER(α/β). In melanoma cells, a two-fold molar excess of G-36 completely blocked G-1 effects (Fig 3F). c-Myc loss following GPER activation was rapid (Fig. 3G) and PKA dependent (Fig. 3H), suggesting that canonical stimulatory G Protein-Coupled Receptor signaling destabilized c-Myc protein. Consistent with this, c-Myc loss after GPER activation was proteasome dependent (Fig. 3I), and c-Myc protein half-life was markedly shortened (Fig. 3J). Together, these data indicate that GPER activation regulates c-Myc through protein degradation.

**Figure 3.**
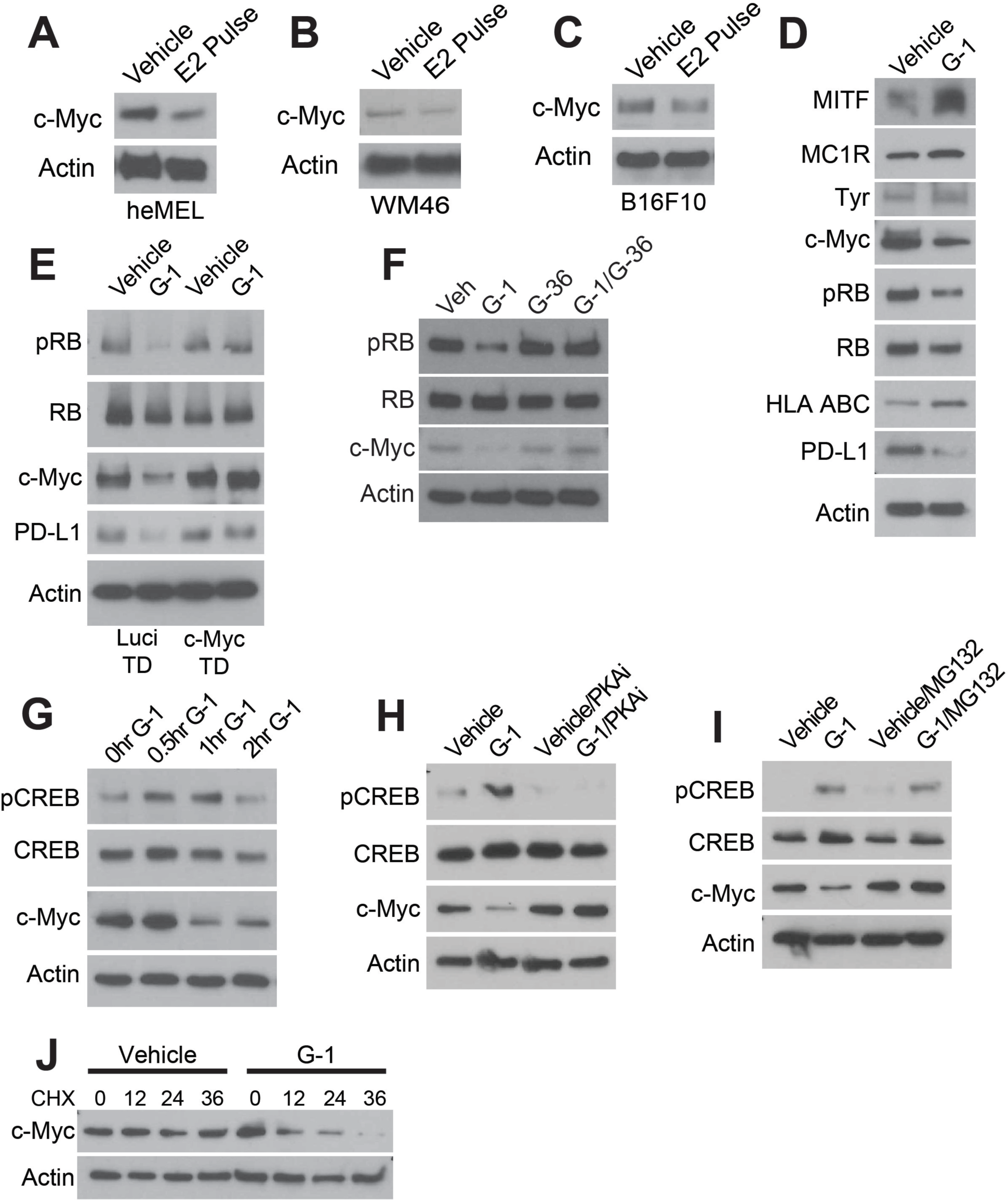
GPER signaling results in loss of c-Myc in melanoma. **A-C**, Western blots of heMel (**A**), WM46 (**B**), and B16F10 (**C**) melanoma transiently treated with E2 for 3 days, followed by 4 day withdraw. **D**, Western blot of WM46 cells treated with a specific GPER agonist (G-1) for 16 hours. **E**, Western blot of luciferase- or c-Myc-transduced WM46 cells treated with G-1 for 16 hours. **F**, Western blot of WM46 cells treated with G-1, GPER antagonist G-36, or a combation for 16 hours. **G**, Western blot of WM46 cells treated with G-1 across a time course. **H**, Western blot of WM46 cells treated with G-1, 100μM PKA inhibitor Rp-8-Br-cAMPS (PKAi), or both for 1 hour. **I**, Western blot of WM46 cells treated with G-1, 2.5μM proteasome inhibitor (MG132), or both for 1 hour. **J**, Western blot of WM46 cells treated with 10μg/ml cyclohexamide (CHX) with and without G-1.

Beyond its role in stimulating proliferation and inhibiting differentiation, c-Myc was recently shown to contribute to tumor aggressiveness by promoting expression of multiple inhibitory immune checkpoint regulators on tumor cells including PD-L1 (23). Consistent with this, pharmacologic GPER activation in melanoma cells resulted in parallel decreases in both c-Myc and PD-L1 (Fig. 4A-C). This PD-L1 depletion was depend on c-Myc loss, as PD-L1 was preserved in cells engineered to maintain normal c-Myc levels in the presence of GPER agonist (Fig. 3E). Given that GPER signaling induced stable changes in tumor cells that antagonized tumor proliferation and decreased tumor cell expression of immune suppressive proteins, we next questioned whether GPER activation potentiates the anti-tumor activity of immune checkpoint blockade inhibitors which are currently the standard of care for advanced melanoma in people (1, 2).

**Figure 4.**
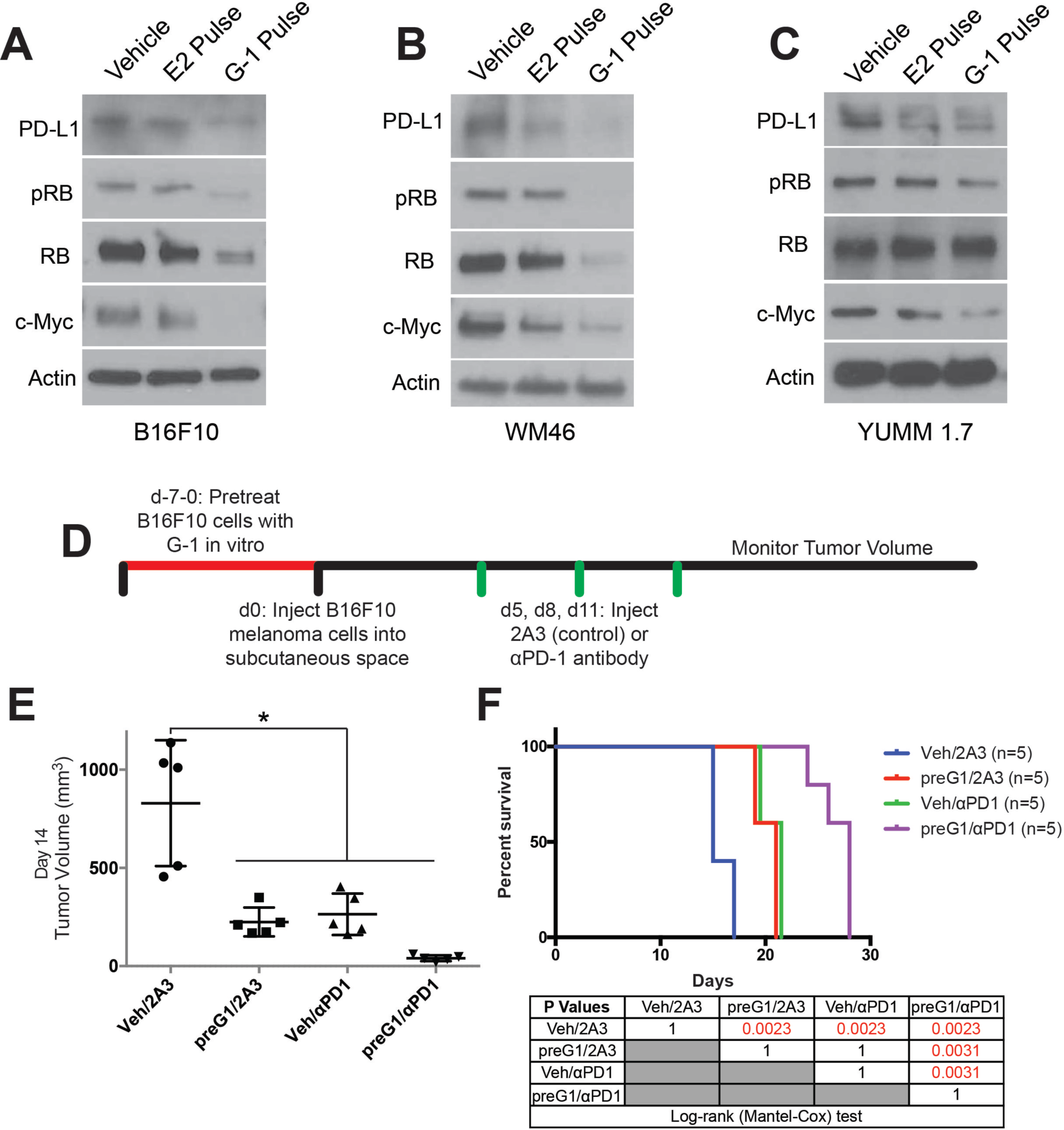
Transient GPER activation inhibits proliferation and augments response to immunotherapy. **A-C**, Western blots of B16F10 (**A**), WM46 (**B**), and YUMM 1.7 (**C**) melanoma cells after transient treatment with a pregnancy-associated concentration of E2 (25 nM) or an optimized concentration of G-1 (500 nM). **D**, Experimental timeline of vehicle or G-1 pre-treatment of B16F10 cells followed by treatment with either αPD-1 antibody or isotype antibody control (2A3), n=5 per group. **E**, Tumor volumes of treatment groups at Day 14, * denotes significance One-way ANOVA with Tukey’s multiple comparison test. **F**, Survival curve of mice with tumors pre-treated with vehicle or G-1, followed by isotype antibody control (2A3) or αPD-1 antibody. Significance between groups by the Log-Rank (Mantel-Cox) test is listed in the table below.

To determine whether tumor cell intrinsic GPER signaling influences melanoma vulnerability to immune checkpoint therapy, we again took advantage of the fact that GPER driven differentiation is long-lasting. We used G-1 to activate GPER and drive differentiation in murine B16F10 melanoma cells in vitro (Fig. 4D). We then injected equal numbers of vehicle or G-1 treated tumor cells into syngeneic C57BL/6 mice, and treated the animals with either αPD-1 antibody or isotype antibody control. Consistent with the persistently differentiated state induced by transient GPER activation, G-1 pretreatment alone inhibited subsequent tumor growth and extended survival compared to controls. αPD-1 antibody monotherapy in animals injected with vehicle treated B16F10 cells also similarly prolonged survival. However, combination of G-1 pretreatment with αPD-1 antibody dramatically extended survival beyond that seen with either agent alone, indicating that GPER activity in tumor cells induced persistent changes in the tumor sufficient to improve the anti-tumor activity of systemically administered αPD-1 therapy (Fig. 4E-F).

To determine whether G-1 may have therapeutic utility as a systemically delivered agent for established melanoma, with or without immune checkpoint inhibitors, mice harboring syngeneic melanoma initiated from untreated B16F10 cells were treated with subcutaneous G-1, αPD-1 antibody, or both, and survival compared to matched mice treated with vehicle and isotype antibody controls (Fig. 5A). G-1 was well tolerated in mice, and G-1 monotherapy extended survival to the same extent as αPD-1. Strikingly, combined treatment with αPD-1 and G-1 extended survival 7 times longer than with either agent alone, indicating a marked synergistic benefit (Fig. 5B-C). Although B16F10 melanoma is the most commonly used model for melanoma immunology studies, and experimental results have largely translated to humans (24, 25), B16F10 lacks the BRaf or NRas oncodriver mutations present in most human melanomas (26, 27). To test whether GPER signaling has similar anti-melanoma activity in a potentially more medically relevant model, we used genetically-defined melanoma cells from the newly-available Yale University Mouse Melanoma collection (YUMM). This resource contains melanoma lines generated from established genetically engineered mouse models that were backcrossed onto C57BL/6 backgrounds specifically to facilitate immunology studies (28). We injected YUMM 1.7 cells (BRaf^V600E/wt^ Pten^−/-^ Cdkn2^−/-^) into C57BL/6 mice, and initiated G-1 treatment with and without αPD-1 after tumors reached 3-4 mm in diameter (Fig. 5D). Similar to results observed with B16F10 melanoma, G-1 or αPD-1 monotherapy significantly extended survival, while combination treatment dramatically extended survival further, including long term survivors (Fig. 5E-F). These results indicate that GPER anti-tumor activity is independent of tumor oncodriver. Consistent with the hypothesis that GPER activation changes the nature of immune infiltration, G-1 treatment in melanoma bearing mice increased several immune cell subsets, including T cells and NK cells, suggesting a more robust inflammatory response (Supplementary Fig. S3A-C).

**Figure 5.**
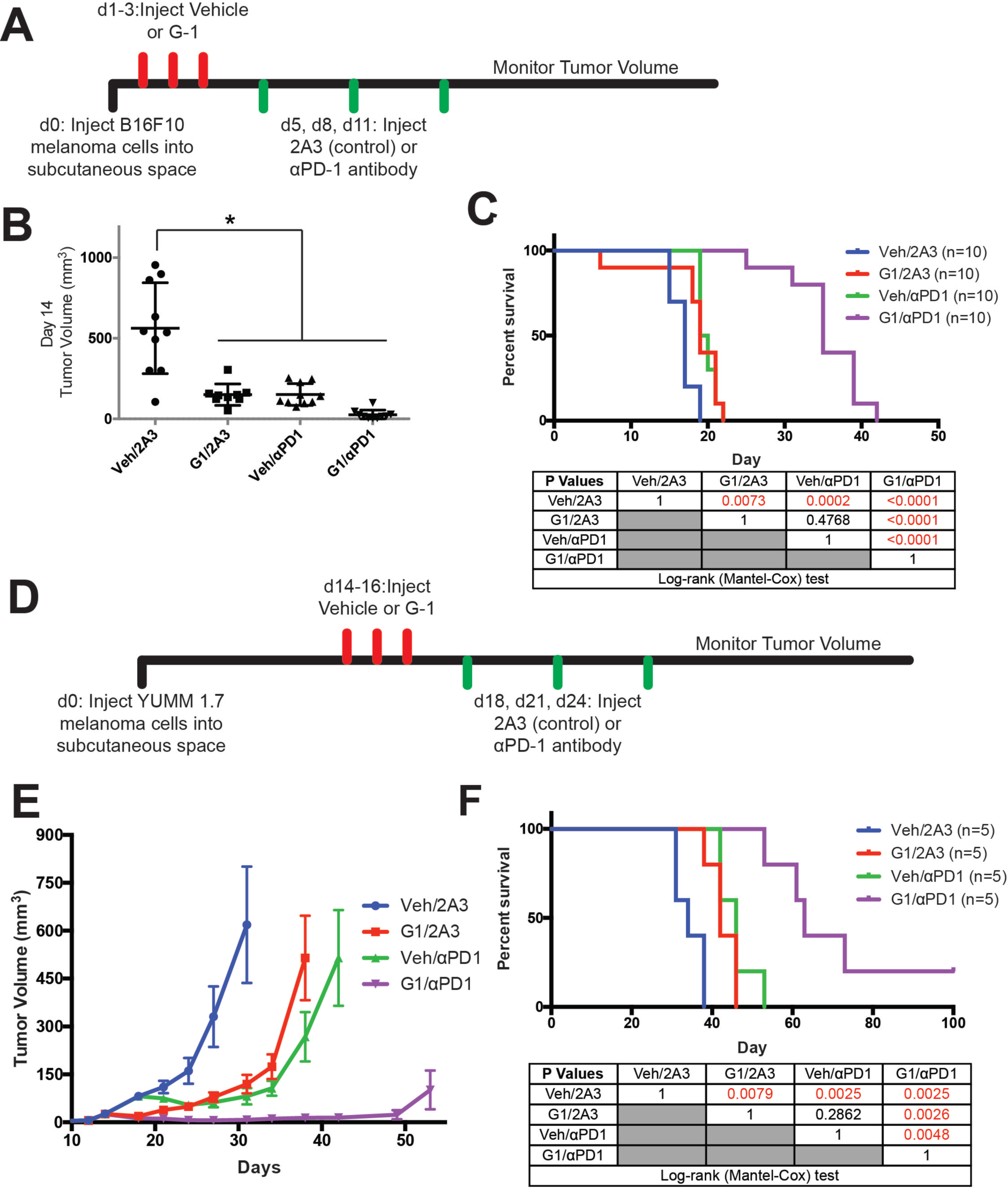
Treatment of melanoma bearing mice with G-1 and αPD-1 immunotherapy dramatically extends survival. **A**, Experimental timeline of B16F10 bearing mice treated with vehicle or G-1, as well as αPD-1 antibody or isotype antibody control (2A3), n = 10 per group. **B**, Tumor volumes of treatment groups at Day 14, * denotes significance One-way ANOVA with Tukey’s multiple comparison test. **C**, Survival curve of mice treated with vehicle or G-1, as well as isotype antibody control (2A3) or αPD-1 antibody. Significance between groups by the Log-Rank (Mantel-Cox) test is listed in the table below. **D**, Experimental outline of YUMM1.7 bearing mice treated with vehicle or G-1, as well as isotype antibody control (2A3) or αPD-1 antibody. Treatment was started at day 14 after tumors reached 4-5 mm in diameter. n = 5 per group. **E**, Tumor volumes over time of treatment groups. **F**, Survival curve of mice treated with vehicle or G-1, as well as αPD-1 antibody or isotype antibody control (2A3). Significance between groups by the Log-Rank (Mantel-Cox) test is listed in the table below.

## Discussion

Although five decades of clinical experience strongly suggest that female sex hormones protect against melanoma, the mechanisms through which pregnancy, or estrogen, influences melanoma have gone relatively unexplored. A pharmacologic approach that recapitulates the female/pregnancy protective effects in men, and women who have not been pregnant, would significantly diminish the overall melanoma burden. However, we are unaware of any previous efforts to harness the melanoma-protective effects of pregnancy for therapeutic benefit. Progress in this area has likely been limited by the fact that estrogen effects in melanocytes are not mediated by the well-known nuclear estrogen receptor, but rather through the nonclassical G protein coupled receptor GPER (8). Here we demonstrate that this nonclassical estrogen signaling promotes differentiation in melanoma, inhibits tumor cell proliferation, and critically, promotes a phenotype that renders tumors more susceptible to immune-mediated elimination. Consistent with this, recent independent work from others has demonstrated that GPER protein levels are higher in human pregnancy-associated melanoma tissue compared to melanomas from non-pregnant females or men, and high GPER expression is associated with favorable prognostic indicators including decreased Breslow depth, decreased mitotic rate, and increased lymphocyte infiltration into tumor (29).

We determined that one of the major mechanisms through which GPER signaling antagonizes melanoma is thorough depletion of c-Myc protein. Although c-Myc amplification is among the most common genetic events in human cancer, and is thus an attractive biologic target, efforts to inhibit Myc with systemically-tolerated agents, have generally been unsuccessful. High c-Myc protein in tumor cells inhibits expression of antigen presenting HLA/MHC (21) and activates expression of PD-L1 (23), which then renders tumors less visible to immune cells. Consistent with this, GPER-induced Myc depletion was accompanied by a reciprocal increases in HLA/MHC protein and decrease in PD-L1 (Fig. 3D), and susceptibility to immune checkpoint inhibitor therapy. As there are currently no FDA approved drugs that directly target Myc, GPER may present a major new therapeutic opportunity.

To our knowledge, this is the first work to demonstrate the potential therapeutic utility of combining differentiation-based therapy with cancer immunotherapy for any cancer type. This approach may also prove useful for other cancers. Differentiation drivers likely have very large “therapeutic windows” as anti-cancer agents. Almost all cancer therapeutics, including immunotherapeutics, are *inhibitors* of oncoproteins, or associated signaling pathways that are also utilized by normal cells. As mammalian evolution has not selected for the ability to compensate for nonphysiologic, pharmacologic inhibition of critical signaling pathways targeted, the utility of most anti-cancer agents is limited by their toxicity to normal tissue. In contrast, we propose targeting GPER with *agonists*. Melanocytes (and other GPER expressing cells) normally respond to physiologic GPER activation, whose natural ligand is endogenous estrogen, and the synthetic specific GPER agonist G-1 is well tolerated in mice. Although no approved drugs target GPER, G Protein-Coupled Receptors are important biologically, and are generally highly “drugable”, as up to 40% of all FDA approved medications act through GPCRs. The specific GPER agonist used here (G-1) may be useful clinically as the first example of a new class of targeted pharmacologics for melanoma.

## Materials and Methods

### Cell culture and cell lines

Primary human melanocytes were extracted from fresh discarded human foreskin and surgical specimens as previously described (14) with some modifications detailed as follows. After overnight incubation in Dispase, the epidermis was separated from the dermis and treated with trypsin for 10 min. Cells were pelleted and plated in selective melanocyte Medium 254 (Invitrogen, Carlsbad, CA, USA) with Human Melanocyte Growth Supplement, and 1% penicillin and streptomycin (Invitrogen). B16F10 melanoma cells were a gift from Andy Minn (University of Pennsylvania Institute, Philadelphia, PA, USA). WM46 melanoma cells were a gift from Meenhard Herlyn (Wistar Institute, Philadelphia, PA, USA). YUMM1.7 melanoma cells were a gift from Ashani Weeraratna (Wistar Institute, Philadelphia, PA, USA) and Marcus Bosenberg (Yale University, New Haven, CT, USA). These cell lines were verified to be of melanocyte origin by response to alpha melanocyte stimulating hormone and melanin production. Human-engineered melanoma cells (heMel) were cultured in Medium 254, WM46 cells were cultured in TU2% media, B16F10 and YUMM1.7 cells were cultured in DMEM (Mediatech, Manassas, VA, USA) with 5% FBS (Invitrogen) and 1% antibiotic-antimycotic (Invitrogen). Cells were transduced with lentiviruses as described previously13. Progesterone (P8783) and 17β-Estradiol (E8875) were purchased from Sigma-Aldrich (St. Louis, MO, USA). G-1 (10008933) and G-36 (14397) were purchased from Cayman Chemical (Ann Arbor, MI, USA). Rp-8-Br-cAMPS was purchased from Santa Cruz Technologies (Dallas, Texas, USA). These compounds were diluted to working stock solutions in Medium 254.

### Mice

All mice were purchased from Taconic (Hudson, NY, USA). Five to seven week old female immune deficient (ICR SCID) and syngeneic (C57BL/6NTac) mice were allowed to acclimatize for one week prior to being used for experiments. These studies were preformed without inclusion/exclusion criteria or blinding, but included randomization. Based on a twofold-anticipated effect, we performed experiments with at least 5 biological replicates. All procedures were performed in accordance with International Animal Care and Use Committee (IACUC)-approved protocols at the University of Pennsylvania.

### Human-engineered melanoma xenografts

Organotypic skin grafts were established using modifications to previously detailed methods (14). The Keratinocyte Growth Media (KGM) used for keratinocyte-only skin grafts was replaced with modified Melanocyte Xenograft Seeding Media (MXSM). MXSM is a 1:1 mixture of KGM, lacking cholera toxin, and Keratinocyte Media 50/50 (Gibco) containing 2% FBS, 1.2 mM calcium chloride, 100 nM Et-3 (endothelin 3), 10 ng/mL rhSCF (recombinant human stem cell factor), and 4.5 ng/mL r-basic FGF (recombinant basic fibroblast growth factor). Briefly, primary human melanocytes were transduced with lentivirus carrying doxycycline-inducible BRAF(V600E), dominant-negative p53(R248W), active CDK4(R24C) and hTERT. Transduced melanocytes (1.5 × 105 cells) and keratinocytes (5.0 × 105 cells) were suspended in 80 µL MXSM, seeded onto the dermis, and incubated at 37°C for 4 days at the air-liquid interface to establish organotypic skin. Organotypic skin tisssues were grafted onto 5-7 week-old female ICR SCID mice (Taconic) according to an IACUC–approved protocol at the University of Pennsylvania. Mice were anesthetized in an isoflurane chamber and murine skin was removed from the upper dorsal region of the mouse. Organotypic human skin was reduced to a uniform 11 mm × 11 mm square and grafted onto the back of the mouse with individual interrupted 6-0 nylon sutures. Mice were dressed with Bactroban ointment, Adaptic, Telfa pad, and Coban wrap. Dressings were removed 2 weeks after grafting and the tissue was allowed to stabilize for an additional week before mice were switched over to doxycycline chow (6g/kg, Bio-Serv, Flemington, NJ) for 15 weeks.

### Subcutaneous tumors and treatments

Subcutaneous tumors were initiated by injecting tumor cells in 50% Matrigel (Corning, Bedford, MA, USA) into the subcutaneous space on the left and right flanks of mice. For each type of tumor injection, 4 × 10^4^ B16F10 cells were used, 1 × 10^6^ WM46 cells were used, and 1 × 10^5^ YUMM1.7 cells were used. In vivo G-1 treatments were performed by first dissolving G-1, synthesized as described previously, in 100% ethanol at a concentration of 1mg/ml. The desired amount of G-1 was then mixed with an appropriate volume of sesame oil, and the ethanol was evaporated off using a Savant Speed Vac (Thermo Fisher, Waltham, MA, USA), leaving the desired amount of G-1 dissolved in 50µL of sesame oil per injection at a 0.4mg/kg dose for B16F10 experiments, and 10mg/kg dose for YUMM1.7 experiments. Vehicle injections were prepared in an identical manner using 100% ethanol. Vehicle and G-1 injections were delivered through subcutaneous injection as indicated in each experimental timeline. Isotype control antibody (Clone: 2A3, BioXcell, West Lebanon, NH, USA) and αPD-1 antibody (Clone: RMP1-14, BioXcell) were diluted in sterile PBS and delivered through intraperitoneal injections at a dose of 10mg/kg.

### Survival Analysis

As subcutaneous tumors grew in mice, perpendicular tumor diameters were measured using calipers. Volume was calculated using the formula L × W^2 × 0.52, where L is the longest dimension and W is the perpendicular dimension. Animals were euthanized when tumors exceeded a protocol-specified size of 15 mm in the longest dimension. Secondary endpoints include severe ulceration, death, and any other condition that falls within the IACUC guidelines for Rodent Tumor and Cancer Models at the University of Pennsylvania.

### Western Blot Analysis

Adherent cells were washed once with DPBS, and lysed with 8M urea containing 50mM NaCl and 50mM Tris-HCl, pH 8.3, 10mM dithiothreitol, 50mM iodoacetamide. Lysates were quantified (Bradford assay), normalized, reduced, and resolved by SDS gel electrophoresis on 4–15% Tris/Glycine gels (Bio-Rad, Hercules, CA, USA). Resolved protein was transferred to PVDF membranes (Millipore, Billerica, MA, USA) using a Semi-Dry Transfer Cell (Bio-Rad), blocked in 5% BSA in TBS-T and probed with primary antibodies recognizing β-Actin (Cell Signaling Technology, #3700, 1:4000, Danvers, MA, USA), BRAF V600E (Spring Bioscience, VE1, 1:500, Pleasanton, CA, USA) c-Myc (Cell Signaling Technology, #5605, 1:1000), CDK4 (Cell Signaling Technology, #12790, 1:1000), p-CREB (Cell Signaling Technology, #9198, 1:1000), CREB (Cell Signaling Technology, #9104, 1:1000), HLA-ABC (Biolegend, w6/32,1:500, San Diego, CA, USA), MC1R (Abcam, EPR6530, 1:1000 Cambridge, MA, USA), p53 (Cell Signaling Technology, #2527, 1:1000), human PD-L1 (Cell Signaling Technology, #13684, 1:1000), mouse PD-L1 (R&D systems, AF1019, 1:500, Minneapolis, MN, USA), p-RB (Cell Signaling Technology, #8516, 1:1000), RB (Cell Signaling Technology, #9313, 1:1000), and tyrosinase (Abcam, T311, 1:1000). After incubation with the appropriate secondary antibody, proteins were detected using either Luminata Crescendo Western HRP Substrate (Millipore) or ECL Western Blotting Analysis System (GE Healthcare, Bensalem, PA).

### Melanin Assay

Cells (1 × 10^5^) cells were seeded uniformly on 6-well tissue culture plates. Cells were treated with vehicle controls, estrogen, or G-1 for 4 days. Cells were then trypsinized, counted, and spun at 300 g for 5 minutes. The resulting cell pellet was solubilized in 120 µL of 1M NaOH, and boiled for 5 min. The optical density of the resulting solution was read at 450 nm using an EMax microplate reader (Molecular Devices, Sunnyvale, CA, USA). The absorbance was normalized to the number of cells in each sample, and relative amounts of melanin were set based on vehicle treated controls.

### Immunohistochemistry and Quantification

Formalin fixed paraffin embedded (FFPE) human skin tissue sections from organotypic tissue was stained for MITF protein expression using a primary antibody to MITF (NCL-L-MITF, Leica Biosystems, Nussloch, Germany), MelanA (NCL-L-MITF, Leica Biosystems), and Ki67 (NCL-L-Ki67-MM1, Leica Biosystems). Staining was performed following the manufacturer protocol for high temperature antigen unmasking technique for immunohistochemical demonstration on paraffin sections. For melanin staining FFPE embedded tissue was subjected to Fontana-Masson stain for melanin as previously described7. Tissue section quantification was performed according to Billings et al. 2015. Briefly, 20X photomicrograph images of representative tissue sections were taken using the Zeiss Axiophot microscope. Tiff files of the images were saved and transferred to Adobe Photoshop where pixels corresponding to MITF or Fontana-Masson staining and epidermal area were selected using the color selection and lasso selection tools. Images corresponding to the single specific color were then analyzed using FIJI (Image J) to determine the number of pixels in each sample and normalized to epidermal area. The numbers of pixels representing Fontana-Masson staining were normalized to the total amount of epidermal counter stain. Ki67 proliferation index was calculated by dividing the number Ki67 positive cells by the total number of MelanA positive cells in the samples.

### Flow Cytometry

Cell surface markers were assessed by incubating single cell suspensions of tissues with primary fluorochrome-labeled antibodies at 4°C for 60 min in PBS with 5% FBS; FITC-anti-mouse-Nkp46 (29A1.4, Biolegend, #137606, 1:50), PE-CF594-anti-mouse-CD8a (53-6.7, BD Pharmingen, #562283, 1:100), PE-Cy5-anti-mouse-CD3ε (145-2C11, Biolegend, #100310, 1:100, PE-Cy7-anti-mouse-I-A/I-E (M5/114.15.2, Biolegend, #107630, 1:600), V450-anti-mouse-CD44 (IM7, Biolegend, #560451, 1:100), AF700-anti-mouse-CD45 (30-F11, Biolegend, #103128, 1:400), APC-Cy7-anti-mouse-F4/80 (BM8, Biolegend, #123118, 1:100), PerCP-Cy5.5-anti-mouse-CD11b (M1/70, BD Pharmingen, #550993, 1:200), BV570-anti-mouse-CD62L (MEL-14, Biolegend, #104433, 1:50), Live/Dead Fixable Aqua Dead Cell Stain Kit, for 405nm excitation (Thermo Fisher Scientific, L-34966, 1:600). Intracellular staining was done using the Fixation/Permeabilization Kit from eBiosciences. Flow cytometric analysis was performed on LSR II Flow Cytometer (BD Biosciences). Collected data were then analyzed using the FlowJo software (Treestar, Ashland, Oregon, USA).

### Statistical Analysis

All statistical analysis was performed using Graphpad Prism 8 (Graphpad Software, La Jolla, CA, USA). No statistical methods were used to predetermine sample size. Details of each statistical test used are included in the figure legends.

## Author Contributions

C.A.N. J.L., B.Z.S., and T.W.R. designed experiments. C.A.N, J.L., J.Z., A.D., B.Z.S., and T.W.R performed experiments. C.A.N. J.L., B.Z.S., and T.W.R. analyzed the data. C.A.N. and T.W.R wrote the manuscript. All authors contributed to editing the manuscript and support the conclusions.

## Acknowledgements

The authors thank the University of Pennsylvania Skin Biology and Disease Research-based center for primary melanocytes and keratinocytes, and for histologic processing and analysis of tissue sections, Andy Minn for B16F10 cells, Meenhard Herlyn for WM46, WM51, WM3702 cells, Ashani Weeraratna and Marcus Bosenberg for YUMM 1.7 cells, and Jeffry Winkler for synthesizing G-1. The authors also thank Sarah Millar, George Cotsarelis, Meenhard Herlyn, John Seykora, David Manning, Pantelis Rompolas and Thomas Leung for critical pre-submission review. T.W.R. is supported by a grant from the NIH/NCI (RO1 CA163566), a Penn/Wistar Institute N.I.H. SPORE (P50CA174523), and the Melanoma Research Alliance. C.A.N was supported by an NIH/NIAMS training grant (T32 AR0007465-32) and an NIH/NCI F31 NRSA Individual Fellowship (F31 CA206325).

## Ethics Statement

This study was performed in strict accordance with the recommendations in the Guide for the Care and Use of Laboratory Animals of the National Institutes of Health. All of the animals were handled according to approved institutional animal care and use committee (IACUC) protocol (#803381) of the University of Pennsylvania.

## Supplementary Figures

**Supplementary Figure 1.**
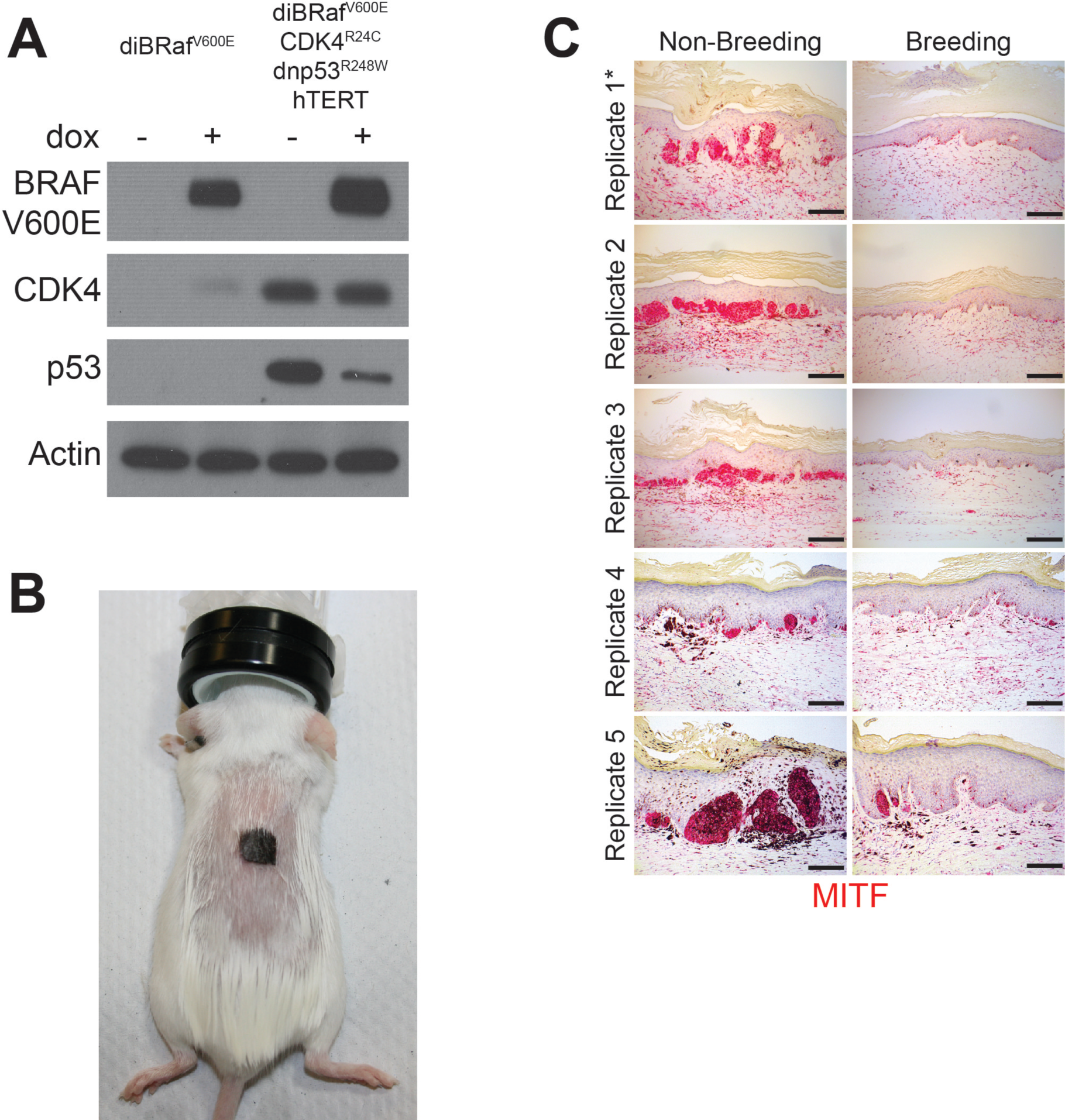
Multiple pregnancies inhibit melanomagenesis. **A**, Western blot validating the transduction of normal human melanocytes with doxycycline inducible BRAF(V600E), dominant-negative p53(R248W), active CDK4(R24C) and hTERT. **B**, Representative photo of a SCID mouse with a human engineered melanoma xenograft. **C**, MITF immunohistochemistry across all non-breeding and breeding mice, * denotes replicates shown in Fig. 1B. Scale bars = 100µM

**Supplementary Figure 2.**
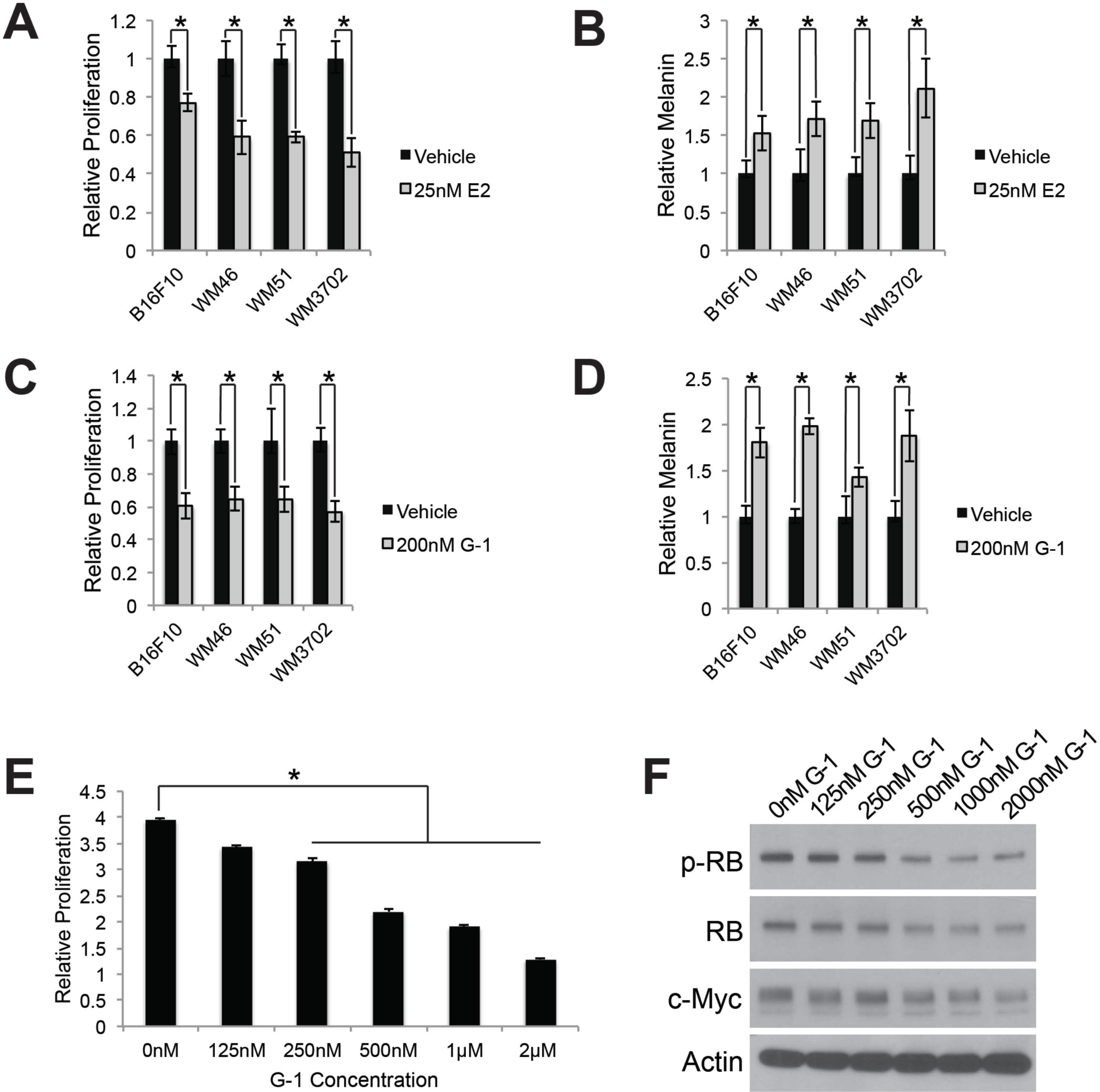
GPER signaling slows proliferation and drives differentiation in mouse and human melanoma. **A**, 5 day proliferation assay of B16F10, WM46 (BRAF^V600E^), WM51 (BRAF^V600E^), and WM3702 (NRAS^Q61L^) cells treated with estrogen (E2), * denotes significance by a two-tailed T-test, n = 3 per group. **B**, 5 day melanin assay of B16F10, WM46 (BRAF^V600E^), WM51 (BRAF^V600E^), and WM3702 (NRAS^Q61L^) cells treated with E2, * denotes significance by a two-tailed T-test, n = 3 per group. **C**, 5 day proliferation assay of B16F10, WM46 (BRAF^V600E^), WM51 (BRAF^V600E^), and WM3702 (NRAS^Q61L^) cells treated with GPER agonist (G-1), * denotes significance by a two-tailed T-test, n = 3 per group. **D**, 5 day melanin assay of B16F10, WM46 (BRAF^V600E^), WM51 (BRAF^V600E^), and WM3702 (NRAS^Q61L^) cells treated with G-1, * denotes significance by a two-tailed T-test, n = 3 per group. **E**, 3 day proliferation assay of B16F10 cells treated with a dose response of G-1, * denotes significance One-way ANOVA with Tukey’s multiple comparison test, n = 5 per group. **F**, Western blot of B16F10 cells treated for 16 hours with a saturating dose response of G-1. All error bars equal the standard deviation of the samples.

**Supplementary Figure 3.**
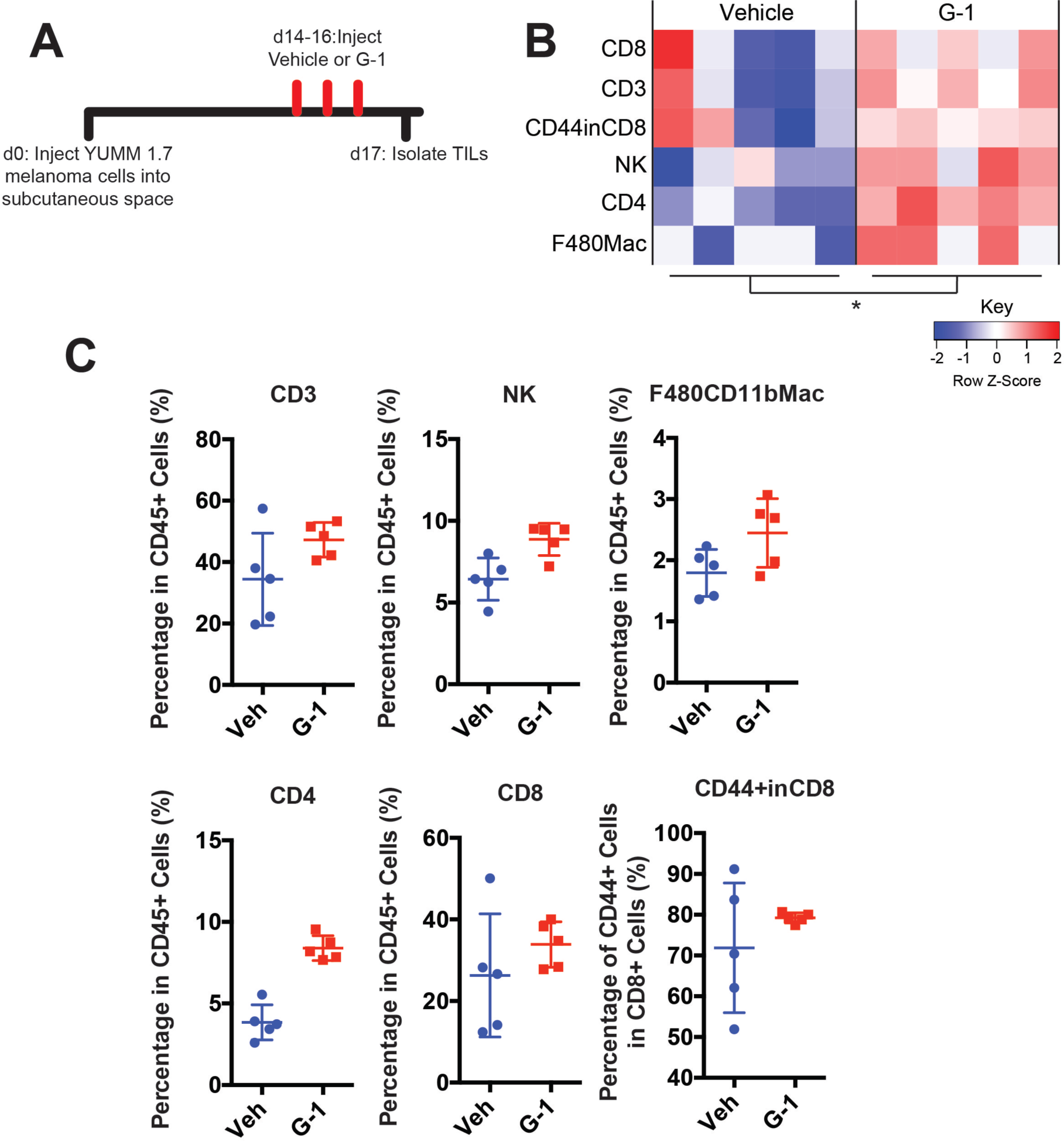
G-1 treatment in vivo alters tumor infiltrating immune cells. **A**, Experimental timeline for vehicle or G-1 treatment of YUMM 1.7 melanoma bearing mice. **B**, Heatmap summarizing immune profiling across biological replicates, n = 5 per group, * denotes significance by two-way ANOVA assuming each immune population is an independent measurement of immune activation. **C**, Quantification of individual immune populations from **B**, n = 5 per group.

